# Subunit Vaccine-Elicited Effector-Like Memory CD8 T Cells Protect Against Listeriosis

**DOI:** 10.1101/2020.12.01.407577

**Authors:** Woojong Lee, Autumn Larsen, Brock Kingstad-Bakke, M. Suresh

**Affiliations:** Department of Pathobiological Sciences, University of Wisconsin-Madison, Madison, 53706, WI, USA

## Abstract

Development of T-cell-based subunit protein vaccines against diseases, such as AIDS, tuberculosis and malaria remains a challenge for immunologists. Here, we have evaluated whether cross-presentation induced by nanoemulsion adjuvant Adjuplex (ADJ), can be combined with the immunomodulatory effects of TLR agonists (CpG or glucopyranosyl lipid adjuvant [GLA]) to evoke protective systemic CD8 T cell-based immunity to *Listeria monocytogenes* (*LM*). Vaccination with ADJ, alone or in combination with CpG or GLA augmented activation and antigen uptake by migratory and resident dendritic cells and up-regulated CD69 expression on B and T lymphocytes in draining lymph nodes. By virtue of its ability to engage BATF3-dependent cross-presenting DCs, ADJ potently elicited effector CD8 T cells that differentiated into a distinct subset of granzyme B-expressing CD27^LO^ effector-like memory CD8 T cells, which provided highly effective immunity to LM in spleen and liver. CpG or GLA alone did not elicit effector-like memory CD8 T cells and induced moderate protection in spleen, but not in the liver. Surprisingly, combining CpG or GLA with ADJ limited the magnitude of ADJ-induced CD8 T cell memory and compromised protective immunity to LM, especially in the liver. Taken together, data presented in this manuscript provides a glimpse of protective CD8 T cell memory differentiation induced by a nano-emulsion adjuvant and demonstrates the unexpected negative effects of TLR signaling on the magnitude of CD8 T cell memory and protective immunity to listeriosis.

**Importance:** To date, the most effective vaccines primarily provide protection by eliciting neutralizing antibodies, while development of T-cell-based subunit vaccines against infectious diseases, such as tuberculosis and malaria, remains a challenge for immunologists. Axiomatically, engagement of multiple innate immune receptors early in the response might be key to programming effective immunity. Hence, there is an impetus to develop combination adjuvants that engage multiple innate signaling pathways and additionally promote cross-presentation to stimulate CD8 T-cell immunity. Here, we show that a nano-emulsion adjuvant ADJ alone elicits effector-like memory CD8 T cells and provides highly effective immunity to listeriosis; combining ADJ with TLR agonists, including CpG and GLA, compromised T cell immunity to LM. In summary, this study provided fundamental insights into the effects of combining innate immune signaling with nano-emulsion adjuvants on memory T cell differentiation and protective immunity. These findings are expected to have implications in the use of combination adjuvants to develop subunit vaccines that engender systemic CD8 T-cell immunity to intracellular pathogens.

## Introduction

Vaccination is a time-tested strategy to control infectious diseases. According to the World Health Organization, there are currently at least 26 vaccines approved for human use. Vaccines are broadly categorized into two types: (A) replicating vaccines (e.g. live attenuated vaccines); (B) non-replicating vaccines (inactivated or subunit vaccines)(1). Live-attenuated vaccines are highly immunogenic and trigger balanced humoral and cell-mediated immunity (2). However, their use is contraindicated in immune-compromised individuals and in pregnancy and there are serious safety concerns regarding adverse events and reversion to virulence (1),(3),(4-6). Highly purified or recombinant subunits of pathogens are poorly immunogenic, which necessitates the use of adjuvants to enhance the immunogenicity of protective antigens in the vaccine (2, 7). Despite decades of vaccine research, very few adjuvants are licensed for use in humans (2, 7-9). Unlike live-attenuated vaccines, current inactivated and subunit vaccines formulated with the licensed adjuvants often confer a shorter duration of immunity, induce antibody biased responses, require multiple immunizations to maintain protective immunity, and trigger poor T_H_1 /CD8 T cell memory (2, 7, 8, 10). A major goal of vaccine development is to identify adjuvants that mimic the immunogenicity and durability of live vaccines. There is emerging consensus that concomitant engagement of multiple innate signaling pathways is a prerequisite to program durable and potent antibody and T cell responses (2).

Adjuvants can be described by two key features: antigen delivery enhancement (e.g. alum, emulsion, liposome) and immune potentiation (Toll-like receptor [TLR] agonists, such as monophosphoryl lipid A [MPL]) (11). Adjuvants and adjuvant combinations consisting of a delivery system and an immune potentiator synergistically enhance antibody and T cell responses (11). T cells have been implicated in protection against varicella, cytomegalovirus, and influenza in humans (2), and there is emerging consensus that protection against infections, such as AIDS, tuberculosis and malaria, requires antibodies, memory CD4 T cells and CD8 T cells (12-15). Therefore, it is critical to identify adjuvant strategies that engender balanced antibody and T cell immunity.

Carbopol polymers (also known as carbomers) are polymers of acrylic acid (16-19) with immune modulating properties (20, 21). Adjuplex (ADJ; Advanced Bioadjuvants) is a nano-emulsion adjuvant composed of carbomers and highly purified soy lecithin. Carbomer-based adjuvants have shown great promise in veterinary vaccines (22, 23), in stimulating neutralizing antibodies against HIV and malaria antigens (24, 25), and also in experimental vaccines against influenza virus in mice (26-29). Combination adjuvants provide effective T cell-based immunity to influenza A virus (29), but is unknown whether vaccines formulated in ADJ can provide T-cell-based protection against systemic infections.

In this study, we tested whether combining ADJ with clinically tested TLR agonists, CpG or glucopyranosyl lipid A (GLA), potentiated the adjuvanticity of ADJ to elicit effective T-cell-based immunity to the intracellular pathogen, *Listeria monocytogenes*. Adjuvants alone or in combination elicited strong innate responses in draining lymph nodes (DLNs), including the activation and engagement of migratory and resident DCs. ADJ effectively activated and expanded antigen-specific effector CD8 T cells to model protein antigen chicken ovalbumin (OVA) by mechanisms dependent upon BATF3-dependent cross-presenting DCs *In vivo*. Notably, effector CD8 T cells elicited by ADJ alone or in combination with CpG or GLA, differentiated into a distinct granzyme B-expressing memory T cell subset termed effector-like memory CD8 T cells. Unexpectedly, combining CpG or GLA with ADJ compromised ADJ-induced protective immunity against listeriosis by limiting the number of memory CD8 T cells and the magnitude of recall T cell responses to listeria challenge. These findings highlight the consequences to the use of combination adjuvants in eliciting effective CD8 T cell-based systemic immunity to intracellular pathogens.

## RESULTS

### Combination adjuvants recruit and activate innate and adaptive cells in vaccine-draining lymph node

We tested whether combining TLR agonists GLA or CpG alone or with adjuplex affected the innate cellular response following vaccination in the footpad. At 24 hours after vaccination, there was a 8- to 10-fold increase in cellularity in popliteal lymph node, the draining lymph node (DLN), in comparison to DLN of mice vaccinated with control PBS or OVA **(Fig. 1A)**. Further analysis of immune cell subsets (**Fig. S1)** revealed a substantive increase in the accumulation of monocytes, neutrophils, XCR1^+^ CD103^+^ migratory DCs, CD8α^+^/CD8α^-^ resident DCs, CD4 T cells, CD8 T cells, and B cells in the DLN **(Fig. 1B),** following vaccination with all adjuvants, as compared to no adjuvant control. There were no significant differences in the number of B or T cells between various adjuvant groups. Notably, GLA alone elicited fewer monocytes, migratory DCs, and lymphoid DCs, in comparison to ADJ, CpG, ADJ+CpG and ADJ+GLA groups. ADJ+CpG recruited the greatest number of neutrophils, among all adjuvants (**Fig. 1B**).

**Figure 1.**
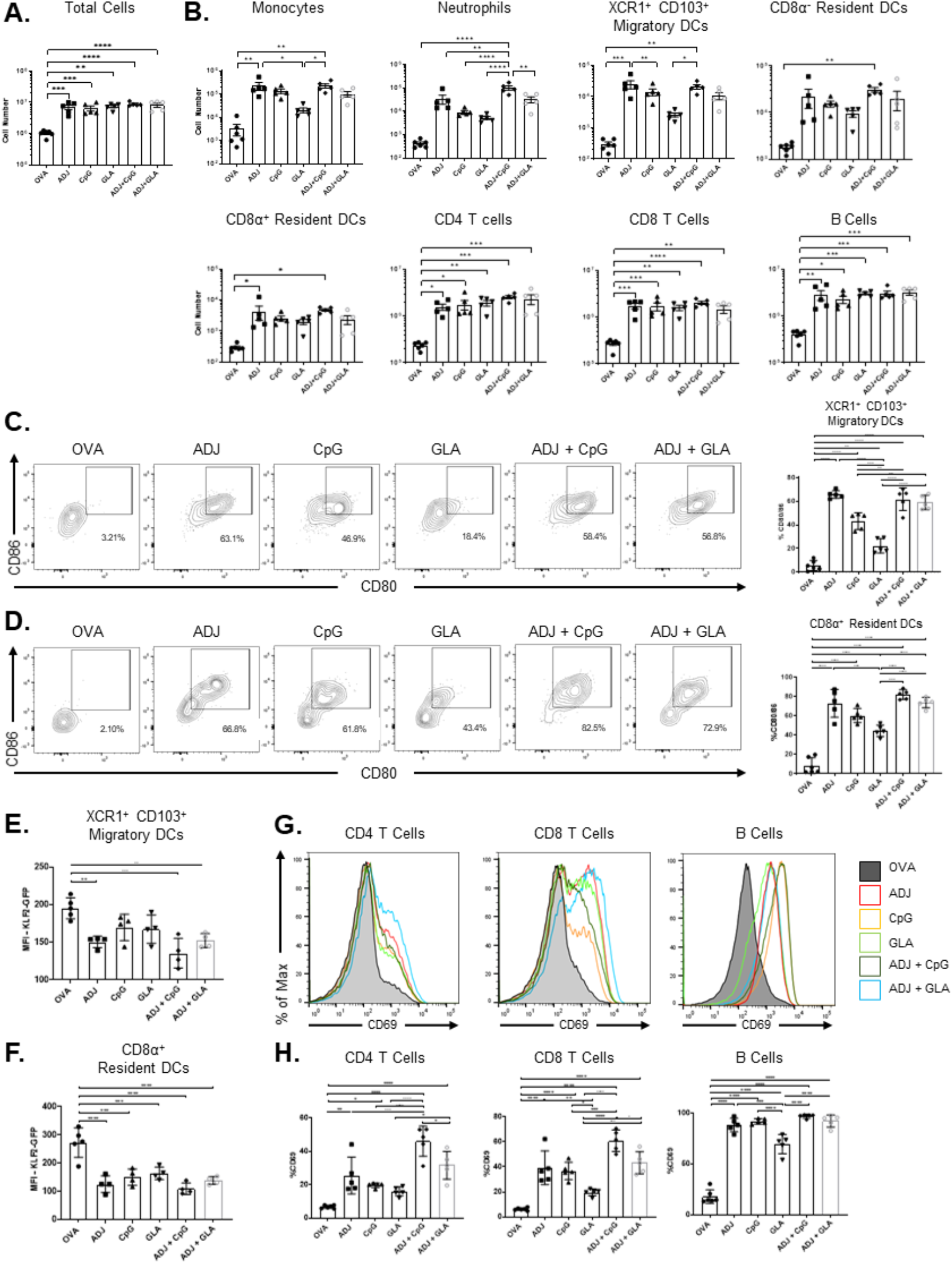
ADJ and TLR-agonist based vaccines induce recruitment and activation of innate and adaptive immune cells in DLNs. Mice were vaccinated SQ with OVA formulated in ADJ, CpG, GLA, ADJ+CpG or ADJ+GLA. Twenty-four hours after vaccination, DLNs were processed into single-cell suspensions and stained with antibodies conjugated to fluorophores. (A) Total cell count of DLN from vaccinated mice. (B) Effect of vaccinations on numbers of Ly6C^+ve^ monocytes, Ly6G^+ve^ neutrophils, XCR1^+ve^ CD103^+ve^ migratory DCs, B220^+ve^ CD19^+ve^ B cells, CD4^+ve^ T cells, and CD8^+ve^ T cells, as assessed by FACS analysis using gating strategy shown in Fig S1. (C-D). Activation status of XCR1^+ve^ CD103^+ve^ migratory DCs (C) and CD8α^+ve^ resident DCs (D), as measured by CD80/86 expression. (E-F) Median fluorescence intensity of KLF2-GFP in XCR1^+ve^ CD103^+ve^ migratory DCs (E) and CD8α^+ve^ resident DCs (F). (G-H) CD69 expression by CD4, CD8, and B cells. Data are the Mean ± SEM from one of two independent experiments with 4-5 mice per group. *, **, ***, and **** indicate significance at *P*<0.1, 0.01, 0.001, and 0.0001 respectively. (One-way ANOVA: A-H)

To further assess the immunostimulatory effects of various adjuvants on DCs, we measured cell surface expression of co-stimulatory molecules CD80 and CD86. At 24 hours post-vaccination, XCR1^+^ CD103^+^ migratory DCs (mDCs) and CD8a^+^ tissue-resident DCs (rDCs) in DLNs of all vaccinated mice displayed significantly (P<0.05) increased expression of CD80 and CD86 **(Fig. 1C-D)**, but ADJ appears to be the strongest activator of CD80/CD86 levels in mDCs and rDCs. Exposure to an inflammatory milieu is known to downregulate the expression of transcription factor Kruppel-like factor 2 (KLF2) in innate myeloid cells (30). Therefore, as a measure of the effect of vaccine-elicited inflammatory response on DCs, we quantified KLF2 expression levels in innate immune cell populations using KLF2-GFP reporter mice. We found that KLF2 expression in XCR1^+^ CD103+ migratory DCs was strongly downregulated by ADJ, as compared to CpG and GLA **(Fig. 1E)**. However, KLF2 downregulation in CD8a^+^ resident DCs was uniformly induced by all adjuvants **(Fig. 1F)**.

Next, we measured the effect of adjuvants on CD4 and CD8 T cells, and B cells in DLNs by measuring the expression of the early activation marker CD69. Within 24 hours of vaccination, all adjuvants triggered higher levels of CD69 expression on B cells, and CD4/CD8 T cells in DLNs, as compared to OVA group **(Fig. 1G-H)**. Note the additive effect of combining ADJ and CpG in inducing CD69 expression on T cells. Together, our data strongly suggested that ADJ, CpG, and GLA effectively recruit and/or activate conventional DCs and T cells in DLN*s*.

### ADJ and TLR agonists enhance antigen uptake by various innate immune cells in vivo

We explored whether various adjuvants affected antigen uptake and activation of APCs in DLNs by vaccinating mice with Alexa Fluor 647-labeled OVA formulated with various adjuvants. At 24 hours after vaccination, all adjuvants significantly augmented the number of OVA-647^+^ cells in DLNs **(Fig. 2A-B)**. We also found that innate immune cells, including monocytes, neutrophils, XCR1^+^ CD103^+^ mDCs and CD8a^+^ rDCs, internalized antigen in vaccinated mice **(Fig. 2C)**. Taken together, data in **Fig. 1 and 2** suggested that ADJ and TLR-based adjuvants augmented varying levels of leukocyte recruitment, DCs’ activation and antigen-containing DCs in DLN.

**Figure 2.**
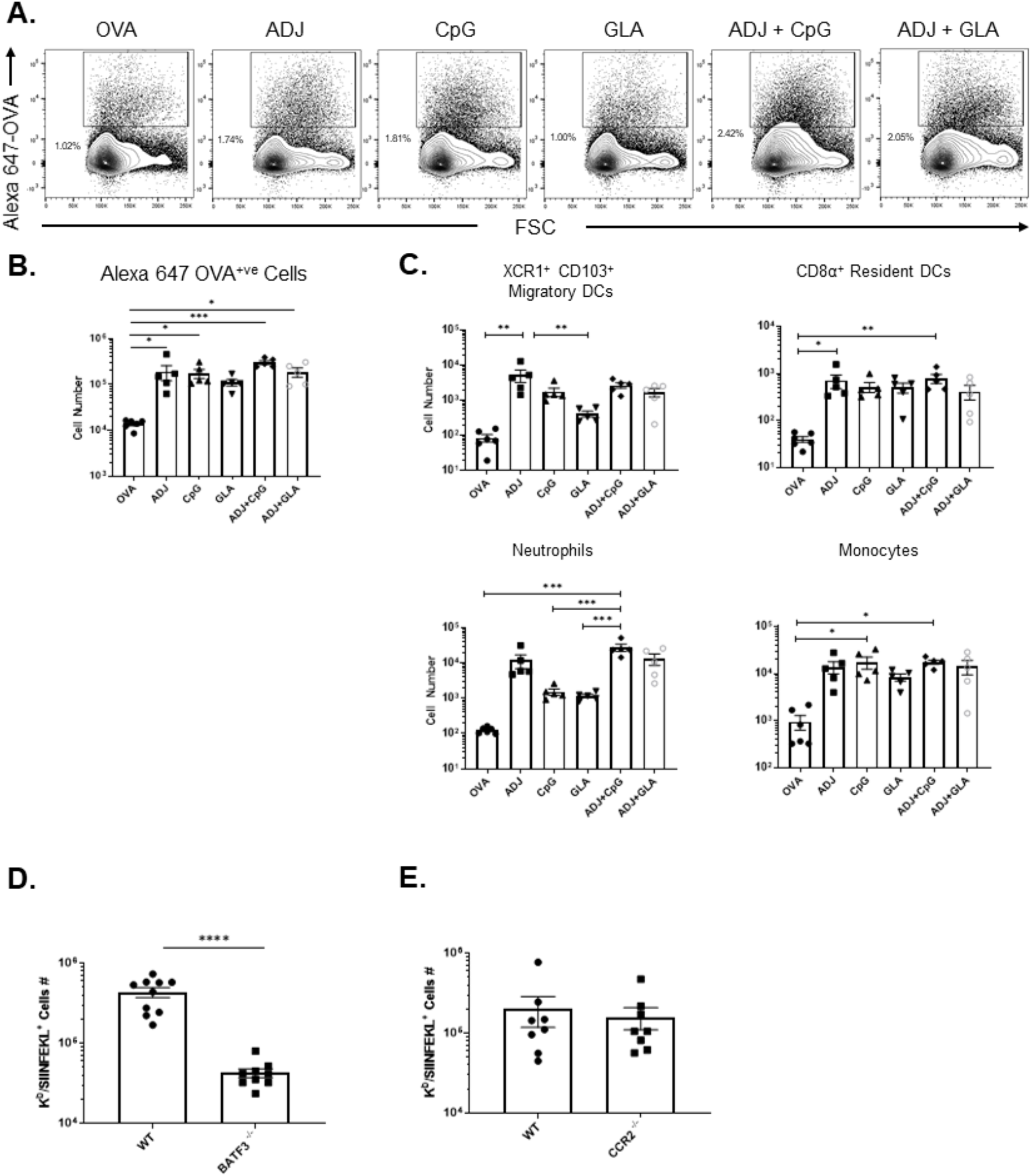
Vaccine adjuvants target antigens to conventional DCs in the DLNs, leading to efficient cross-presentation and CD8 T cell activation *in vivo*. Mice were vaccinated SQ with Alexa Fluor 647 (AF 647)-conjugated OVA formulated in ADJ, CpG, GLA, ADJ+CpG, or ADJ+GLA. Twenty-four hours after vaccination, DLNs were mechanically processed into single-cell suspensions. (A) Antigen-containing cells were visualized using AF647-conjugated OVA by flow cytometry. (B) Total numbers of AF647-OVA^+ve^ cells. (C) Total numbers of AF647-OVA+ immune cell subsets (XCR1^+ve^ CD103^+ve^ migratory DCs, CD8α^+ve^ resident DCs, neutrophils, and monocytes). (D) Wild type (WT) B6 and BATF3-deficient (BATF3-/-) mice were vaccinated SQ with OVA (10ug) formulated in ADJ (5%). On the 8th day after vaccination, the total number of activated OVA SIINFEKL-specific CD8 (CD44^+ve^, K^b^/SIINFEKL^+ve^) T cells in the spleen were quantified by flow cytometry. (E) WT B6 and CCR2-deficient (CCR2-/-) mice were vaccinated SQ with OVA (10ug) + ADJ (5%). On the 8th day after vaccination, the total number of activated OVA SIINFEKL-specific CD8 (CD44^+ve^, K^b^/SIINFEKL^+ve^) T cells in the spleen were quantified by flow cytometry. Data are Mean ± SEM; the data represent one of two independent experiments (A-C) or are pooled from two independent experiments (D-E). *, **, ***, and **** indicate significance at *P*<0.1, 0.01, 0.001, and 0.0001 respectively. (One-way ANOVA: A-C; Mann-Whitney U test-D-E).

### Effect of TLR agonists on ADJ-driven DC cross-presentation in vitro and the role of BATF3 on ADJ-induced CD8 T cell responses in vivo

We have previously demonstrated that ADJ enhanced antigen cross-presentation by DC-like cells *in vitro* (28); it was of interest to investigate whether inclusion of TLR agonists synergistically augmented ADJ-induced DC cross-presentation *in vitro*. To this end, BMDCs were treated with OVA formulated in ADJ, CpG, GLA, ADJ+CpG or ADJ+GLA. Subsequently, BMDCs were evaluated for their capacity to activate SIINFEKL-specific B3Z T cell hybridoma cells using a reporter assay (31, 32). ADJ-treated BMDCs potently activated β-gal production in B3Z cells, whereas BMDCs treated with CpG or GLA failed to activate B3Z cells to levels greater than in OVA-stimulated BMDCs **(Fig. S2)**. Cross-presentation by BMDCs treated with ADJ+CpG or ADJ+GLA was comparable to that in ADJ-treated BMDCs. Hence, TLR agonists failed to further augment ADJ-induced cross-presentation by DCs, *in vitro*.

BATF3-dependent DCs are required for effective cross-presentation and activation of CD8 T cells *in vivo* (33). To evaluate the requirement for BATF3-dependent DCs in cross-priming CD8 T cells by ADJ, we immunized wild type (WT) and BATF3-deficient (BATF^-/-^) mice subcutaneously (SQ) with OVA formulated in ADJ. At day 8 after vaccination, we enumerated OVA SIINFEKL-specific CD8 T cells in spleen. We found that BATF3 deficiency abolished the activation and expansion of SIINFEKL-specific CD8 T cells in spleen, which suggested that BATF3 is required for ADJ-driven CD8 T cell responses to subunit vaccines **(Figure 2D)**.

Because we have observed increased numbers of antigen-containing monocytes in DLNs of mice vaccinated with various adjuvants, we next interrogated whether monocytes are required in cross-priming CD8 T cells by ADJ-based subunit protein vaccines. To this end, we vaccinated cohorts of WT and CCR2-deficient (CCR2-/-) mice SQ with ADJ+OVA, and quantified OVA SIINFEKL-specific CD8 T cells in spleens at day 8 after immunization. CCR2 deficiency did not significantly affect the accumulation of SIINFEKL-specific CD8 T cells in spleens, suggesting that monocytes were not required for ADJ-driven antigen cross-presentation and/or expansion of CD8 T cells *in vivo* **(Figure 2E)**.

### Differentiation of effector CD8 T cells in spleen following vaccination with ADJ/TLR-agonist-based combination adjuvants

To investigate whether combination adjuvants differed in terms of the magnitude and nature of the CD8 T cell response, we vaccinated mice SQ twice (21 days apart) with OVA formulated in ADJ, CpG, GLA, ADJ+CpG or ADJ+GLA. At day 8 after boost, we quantified OVA SIINFEKL-specific CD8 T cells in spleens **(Fig. 3A)**. Frequencies and numbers of SIINFEKL-specific CD8 T cells in spleen of ADJ groups were significantly (P<0.05) higher than in CpG or GLA groups. Notably, CpG and GLA failed to further enhance ADJ-induced splenic CD8 T cell responses.

**Figure 3.**
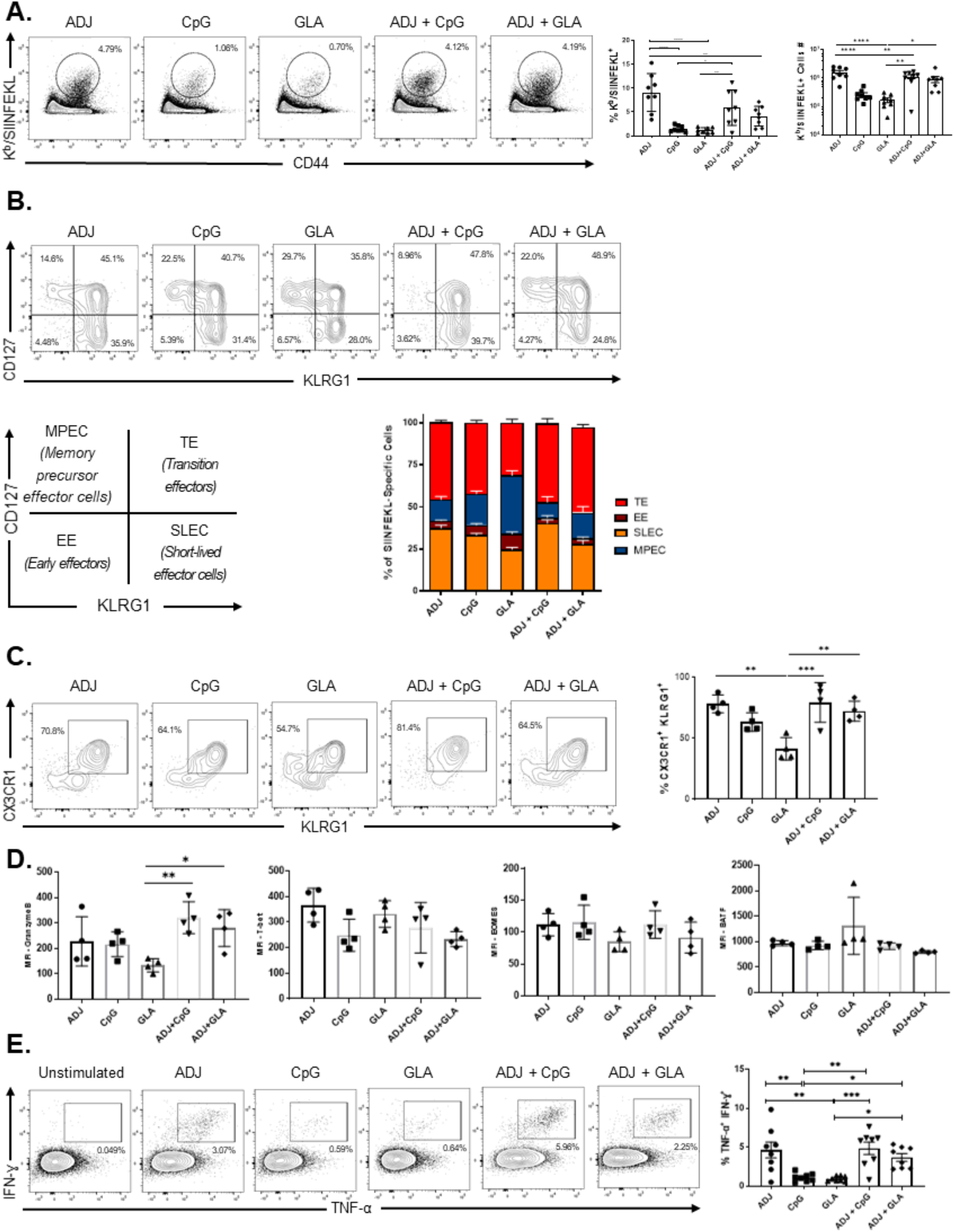
Effector CD8 T-cell responses to adjuvanted vaccines. Mice were vaccinated SQ with OVA formulated in ADJ, CpG, GLA, ADJ+CpG, or ADJ+GLA. Twenty-one days after the first vaccination, mice were boosted with the same formulation. On the 8^th^ day after booster vaccination, single-cell suspensions from spleen were stained with D^b^/SIINFEKL tetramers, anti-CD8, anti-CD44, anti-CD127, anti-KLRG-1, anti-CX3CR1, anti-granzyme B, anti-T-bet, anti-EOMES, and anti-BATF antibodies. (A) Percentages and total numbers of OVA SIINFEKL-specific CD8 T cells in spleens. FACS plots in A are gated on total CD8 T cells and the numbers are the percentages of tetramer-binding cells among the gated population. (B-D) FACS plots are gated on tetramer-binding CD8 T cells and numbers are percentages of the gated cells, in respective gates/quadrants. (E) Median fluorescence intensities (MFI) for transcription factors in SIINFEKL-specific CD8 T cells. (F) Functional polarization of effector CD8 cells. The percentages of CD8 T cells that produced IFN-γ and TNF-α following stimulation with SIINFEKL peptide were quantified by intracellular cytokine staining. Unstimulated control without the SIINFEKL peptide served as a negative control. Data in each graph indicate Mean ± SEM. Data are pooled from two independent experiments (A, E) or represent one of two independent experiments (B-D). Each independent experiment had n=3-5 mice per group. *, **, ***, and **** indicate significance at *P*<0.1, 0.01, 0.001, and 0.0001 respectively. (One-way ANOVA: A-F)

We quantified CD127 and KLRG-1 expression to determine the differentiation state of SIINFEKL-specific effector CD8 T cells as: short-lived effector cells (SLECs; CD127^LO^/KLRG-1^HI^), memory precursor effector cells (MPECs; CD127^HI^/KLRG-1^LO^) and transition effector cells (TEs; CD127^Hi^/ KLRG1^HI^) in spleens of vaccinated mice **(Fig. 3B)**. ADJ and CpG induced the highest levels of KLRG-1 expression, as compared to GLA alone, with no significant differences in CD127 expression between groups. Consequently, the relative proportions of CD127^Hi^/KLRG-1^HI^ TEs were higher in ADJ and CpG groups, as compared to the GLA group. Combining ADJ with CpG reduced the percentages of MPECs, as compared to CpG alone. In comparison to GLA mice, the combination of ADJ and GLA enhanced the percentages of TEs, at the expense of MPECs in the GLA group. Similar to KLRG-1 expression, ADJ was linked to elevated expression of CX3CR1 in SIINFEKL-specific effector CD8 T cells (**Fig. 3C**). The high-level induction of KLRG-1 and CX3CR1 by ADJ was not linked to significant (P<0.05) alterations in the expression of transcription factors T-bet, EOMES or BATF (**Fig. 3D**). In summary, ADJ appeared to promote greater terminal differentiation of effector CD8 T cells in vaccinated mice than other tested adjuvant combinations.

Next, we questioned whether different combinations of adjuvants affected the functionality of effector CD8 T cells. ADJ and TLR agonists induced readily detectable levels of the effector molecule granzyme B, and combining CpG or GLA with ADJ tended to promote granzyme expression, especially compared to GLA group (**Fig. 3D**). SIINFEKL-specific CD8 T cells induced by all adjuvants exhibited polyfunctionality and produced both IFN-γ and TNF-α, upon antigenic stimulation *ex vivo*. The differences in the frequencies of cytokine-producing CD8 T cells among different groups reflect varying frequencies of antigen-specific CD8 T cells (**Fig. 3E**).

### Effector-like CD8 T-cell memory in mice vaccinated with combination adjuvants

40 days after booster vaccination, we quantified and characterized SIINFEKL-specific memory CD8 T cells in spleens. Spleens of mice vaccinated with ADJ alone contained significantly (P<0.05) higher frequencies and numbers of SIINFEKL-specific memory CD8 T cells, than in other groups (**Fig. 4A**). Strikingly, spleens of mice vaccinated with ADJ, ADJ+CpG and ADJ+GLA were enriched for KLRG-1^HI^/CX3CR-1^HI^/CD27^LO^/granzyme B^Hi^ SIINFEKL-specific memory CD8 T cells, that are reminiscent of effector-like memory CD8 T cells described by Jameson’s group (34) (**Fig. 4B-E**). These data suggested that ADJ drove the differentiation of effector-like memory CD8 T cells following vaccination; combining CpG or GLA with ADJ might have dampened the number of memory CD8 T cells in spleen. We next assessed the ability of memory CD8 T cells to produce cytokines, upon *ex vivo* antigenic stimulation. A substantive proportion of CD8 T cells in spleens from the ADJ group produced both IFN-γ and TNF-α, upon stimulation with the SIINFEKL peptide (**Fig. 4F)**. Thus, ADJ promoted the differentiation of highly functional effector-like SIINFEKL-specific memory CD8 T cells in spleens of vaccinated mice.

**Figure 4.**
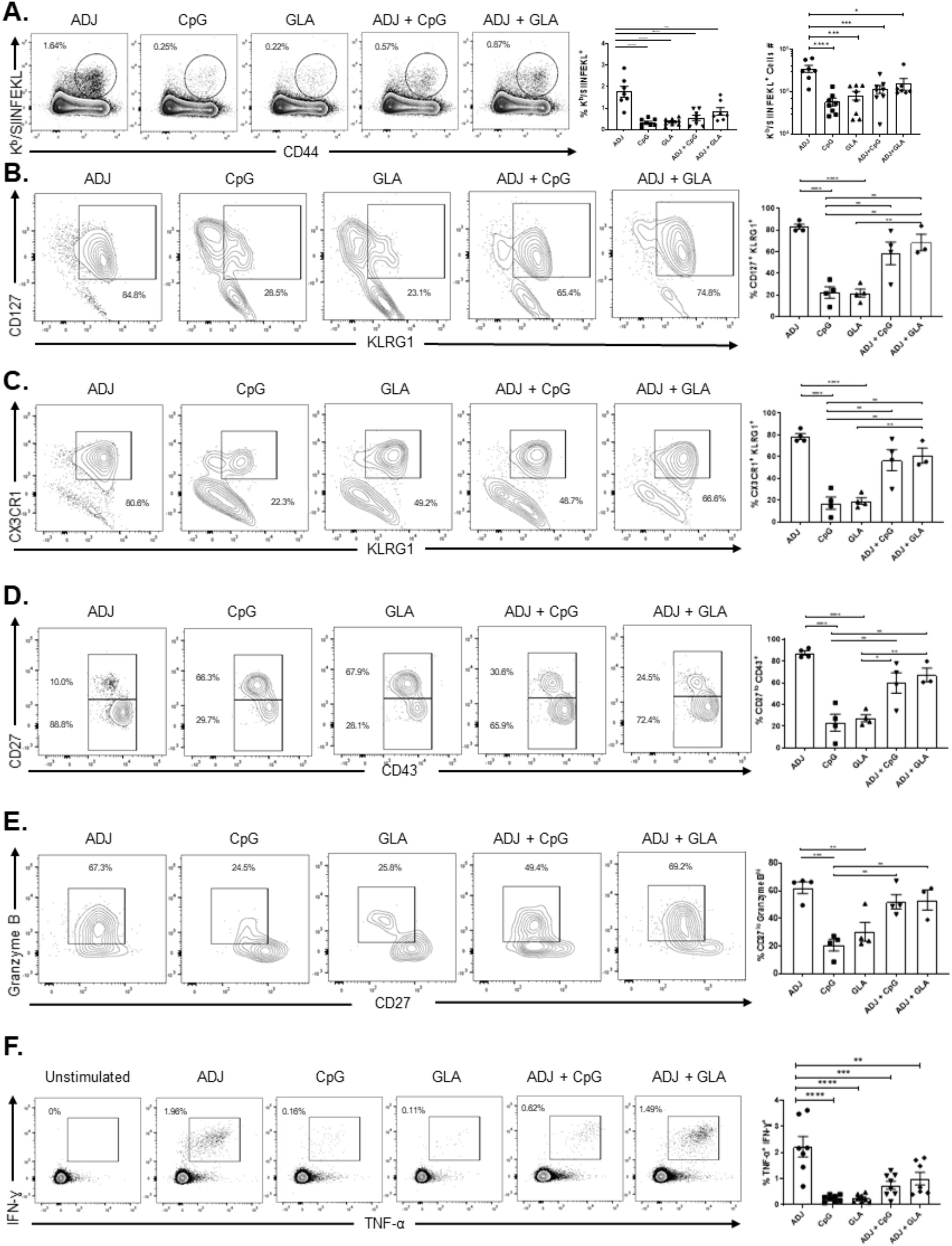
Vaccine-induced CD8 T-cell memory. Mice were vaccinated twice (3 weeks apart) SQ with OVA (10ug/mouse) formulated in ADJ, CpG, GLA, ADJ+CpG, ADJ+GLA. Forty days after the booster vaccination, single-cell suspensions cells of spleen were stained with D^b^/SIINFEKL tetramers, anti-CD8, anti-CD44, anti-CD127, anti-KLRG-1, anti-CX3CR1, anti-CD43, anti-CD27, and anti-granzyme B antibodies. (A) Percentages and total numbers of OVA SIINFEKL-specific CD8 T cells in spleens. (B-E) FACS plots show percentages of gated tetramer-binding CD8 T cells in respective gates/quadrants. (F) Functional polarization of memory CD8 T cells. The percentages of SIIFNEKL-specific CD8 T cells that produced IFN-γ and TNF-α were quantified by intracellular cytokine staining. Cells cultured without SIINFEKL peptide stimulation were used as a negative control. Data in each graph show Mean ± SEM. Data are pooled from two independent experiments (A, F) or represent one of two independent experiments (B-E). Each independent experiment had n=3-5 mice per group. *, **, ***, and **** indicate significance at *P*<0.1, 0.01, 0.001, and 0.0001 respectively. (One-way ANOVA: A-F)

### Addition of TLR agonists compromises ADJ-induced CTL immunity to listeriosis

To assess CD8 T cell memory-dependent protection against listeriosis, we vaccinated mice with OVA formulated in aforementioned adjuvants. Forty days after booster vaccination, mice were challenged with virulent recombinant *Listeria monocytogenes* expressing OVA (LM-OVA). On the 5^th^ day after LM-OVA challenge, we quantified bacterial burden in spleens and liver, and recall T cell responses in spleens. High titers of listeria were detected in the spleens and livers of mice that were unvaccinated or vaccinated with OVA **(Figs. 5A-B)**. Tissues of ADJ-vaccinated mice contained the lowest bacterial burden and provided the most effective protection in spleen and liver, as compared to unvaccinated and other adjuvant groups. Protections afforded by CpG, GLA, ADJ+CpG and ADJ+GLA vaccination were comparable in spleen, but significantly (P< 0.05) better than in OVA group. However, vaccination with CpG, GLA, ADJ+CpG or ADJ+GLA failed to control listeria burden in the liver. Notably, associated with better bacterial control, SIINFEKL-specific memory CD8 T cells in ADJ mice displayed the strongest recall responses and expressed lowest granzyme B levels, as compared to other adjuvant groups (Figs. 5C-F). Thus, combining CpG or GLA with ADJ compromised ADJ-induced T-cell based protective immunity to listeriosis, particularly in the liver.

## Discussion

Establishment of durable and protective memory T cells remains an unrealized goal of developing vaccines against infectious diseases that require T-cell based immunity, including HIV/AIDS, malaria, and tuberculosis (13-15, 35). Studies of live attenuated yellow fever vaccine suggest that engagement of multiple innate immune receptors during the very early phase of the immune response might be essential for programming durable immunity to vaccinations (36, 37). Using ADJ, a polyacrylic acid-based nano-emulsion adjuvant that is known to induce neutralizing antibodies against HIV and malaria (22-25), we have probed whether combining ADJ with TLR agonists enhanced T cell-based vaccine immunity to listeriosis and defined the differentiation state and the phenotypic and functional attributes of antigen-specific memory CD8 T cells induced by combination adjuvants.

It is widely believed that elicitation of less differentiated central memory CD8 T cells that have high proliferative capacities are crucial for enduring T-cell immunity (38). Interestingly, however, Olson *et al*. reported the existence of protective effector-like memory CD8 T cells, which display constitutive cytolytic activity but limited proliferative potential in listeria-immune mice. Such effector-like memory CD8 T cells exhibited phenotypic attributes of terminal differentiation, including high levels of KLRG-1 and diminished levels of CD27, but provided the most effective systemic protection against listeria. The induction of such effector-like memory CD8 T cells by adjuvanted vaccines has not been reported to date. Unexpectedly, we find that ADJ-based vaccines potently induced differentiation of effector-like memory CD8 T cells (KLRG1^HI^/CX3CR1^HI^/CD43^HI^/CD127^HI^/CD27^LO^/Granzyme-B^HI^) in spleen, while GLA or CpG did not. According to the linear model of memory T cell differentiation, activated T cells progress towards terminal differentiation and lose their memory potential as a function of the cumulative strength of antigenic stimulation and the degree of inflammation (38). It is unknown where effector-like memory CD8 T cells emerge from in the spectrum of T cell differentiation. We theorize that effector-like memory CD8 T cells emerge from an intermediate state of differentiation that give to rise to effector memory cells and terminal effector cells. It will be interesting to test whether this distinct hybrid state is a sequel to an epigenetically imprinted constitutive effector program in conjunction with a CD127-driven IL-7-dependent survival program.

How ADJ promotes the differentiation of effector-like memory CD8 T cells remains unknown. Since effector-like memory CD8 T cells are induced during listeriosis (34), we speculate that the immunological milieu and/or the strength of antigenic stimulation experienced by responding CD8 T cells following ADJ-based vaccination mimics listeria infection, leading to the programming of effector-like memory CD8 T cells. All adjuvants induced comparable levels of antigen-containing innate cells *in vivo;* combining CpG or GLA with ADJ did not enhance or negatively affect ADJ-driven DC cross-presentation *in vitro*. However, one of the notable observations was that ADJ drove the highest expression of co-stimulatory molecules, including CD80 and CD86, and induced the most potent inflammatory response, as evidenced by downregulation of KLF2 expression, in XCR1^+^ CD103^+^ mDCs. It is noteworthy that, as compared to CD8α^-^ and CD103^-^ cDC subsets, CD8α^+^ and CD103^+^ cDC subsets are the most prominent cross-presenting DC subsets and immunization studies in Batf3^-/-^ mice strongly suggest an important role for CD8α^+^ rDCs and CD103^+^ mDCs in vaccine-elicited T cell responses (33). Therefore, it is possible that the level of T cell signaling (co-stimulatory and inflammatory) evoked by ADJ (29) is conducive for differentiation of effector-like memory CD8 T cells. More mechanistic studies are warranted to evaluate the nature of the inflammatory milieu and the strength/duration of TCR signaling in regulating the differentiation of effector-like memory CD8 T cells in DLNs.

Parenteral vaccination with ADJ alone provided the most effective protection against listeria challenge, as compared to CpG, GLA, ADJ+GLA and ADJ+CpG. Unexpectedly, combining CpG or GLA with ADJ compromised ADJ-induced protective immunity. The diminished protective immunity induced by combination adjuvants ADJ+GLA or ADJ+CpG was linked to reduced accumulation of effector cells (clonal burst size) and consequent reduction in the number of memory CD8 T cells. Reduced number of memory CD8 T cells, in turn, limited the magnitude of recall T-cell responses and compromised listeria control in mice vaccinated with ADJ+GLA or ADJ+CpG. Less effective listeria control in CpG, GLA, ADJ+CpG and ADJ+GLA was not associated with impaired effector functions, including cytokine production or granzyme B expression of CD8 T cells. Interestingly however, CD8 T-cell expression of granzyme B in listeria-challenged mice showed a clear negative correlation with listeria load in spleen and liver. In the spleen of ADJ mice that effectively controlled listeria, SIINFEKL-specific CD8 T cells expressed lower levels of granzyme-B, as compared to other vaccine groups, especially the CpG and GLA groups. It is likely that higher granzyme B expression in CD8 T cells reflects ongoing antigenic stimulation due to higher bacterial load in mice with compromised vaccine-induced immunity. Although it is unclear how combining CpG or GLA with ADJ negatively modulates the accumulation of effector CD8 T cells and protective immunity, the data presented in this manuscript nevertheless strongly suggest that combination adjuvants may not always enhance vaccine-induced T-cell immunity. It is noteworthy to highlight that unlike parenteral vaccination, intranasal vaccination with combination adjuvants, such as ADJ+GLA and ADJ+CpG, markedly enhance ADJ-induced pulmonary T-cell immunity to influenza A virus (29). Therefore, tenets for inducing systemic versus mucosal T-cell immunity with combination adjuvants are different, and further studies are essential to elucidate the underlying mechanisms.

In summary, we have systematically evaluated the spectrum of effects evoked by a nano-emulsion adjuvant in combination with immunomodulatory adjuvants CpG and GLA in terms of antigen uptake by various innate immune cells, accumulation of innate cells, polyclonal activation of B and T cells, inflammation-induced downregulation of KLF2 in DCs and activation of migratory and lymphoid-resident DCs in DLNs. Furthermore, we document the effect of combination adjuvants on the clonal burst size of effector cells, the magnitude and nature of CD8 T cell memory, the magnitude of recall T cell responses and T cell-based protective immunity to the prototypic intracellular pathogen, *Listeria monocytogenes*. These studies provided unexpected insights into the nature of vaccine-elicited T-cell memory and protective immunity to intracellular pathogens. Further, contrary to prevailing opinion in the vaccine adjuvant field, we found that combining adjuvants might have unintended negative effects on T-cell-based protective immunity. Taken together, findings reported in this manuscript provide new insights into the differentiation of CD8 T cells induced by adjuvanted subunit vaccines, which might pave the way for the rational development of adjuvants that elicit effective T-cell immunity to intracellular pathogens.

## METHODS

### Experimental animals

7-12-week-old C57BL/6J (B6) were purchased from restricted-access SPF mouse breeding colonies at the University of Wisconsin-Madison Breeding Core Facility or from Jackson Laboratory. Batf3^-/-^ (Stock number: 013755) and CCR2^-/-^ (Stock number: 004999) were purchased from Jackson Laboratory. KLF2-GFP reporter mice were provided by Dr. Jameson (University of Minnesota, Minneapolis, MN).

### Ethics statement

All experiments were performed in accordance with the animal protocol (Protocol number V5308 or V5564) approved by the University of Wisconsin School of Veterinary Medicine Institutional Animal Care and Use Committee. The animal committee mandates that institutions and individuals using animals for research, teaching, and/or testing much acknowledge and accept both legal and ethical responsibility for the animals under their care, as specified in the Animal Welfare Act and associated Animal Welfare Regulations and Public Health Service (PHS) Policy.

### B3Z cell hybridoma and BMDC generation

The B3Z hybridoma was a generous gift from Dr. Bruce Klein (University of Wisconsin-Madison). B3Z cells were maintained in Iscove’s Modified Dulbecco’s media supplemented with 10% FBS, 100 U/ml penicillin G and 100 g/ml streptomycin sulfate. Primary cultures of bone marrow-derived DCs were generated as previously described (39, 40).

### B3Z activation assay for *in vitro* cross presentation

The cross-presentation capacity of murine BMDCs was measured using B3Z hybridoma cells, as previously described (31, 32). Briefly, DCs were plated at 1 x 10^5^ cells/well in 96-well round bottom culture-treated plate (Corning). BMDCs were cultured with OVA (1mg/ml) in combination of different adjuvants (ADJ [1%], CpG [5ug/ml], GLA [1ug/ml]) for 5 hours. Next, BMDCs were fixed with 0.025% glutaraldehyde for 2 minutes at room temperature, washed with PBS and cultured with B3Z cells (1 x 10^5^ cells/well) for 18 hours. After 18 hours, B3Z cells were washed and incubated with CPRG substrate (0.15mM, Santa Cruz Biotechnology, sc-257242) in 200ul of lysis buffer (0.1% NP 40+ PBS) for 18 hours at room temperature. The absorbance (590nm) was measured using a plate reader. Wells containing B3Z cells + BMDCs without OVA served as background control.

### Immunization

Hen egg white ovalbumin grade-V (OVA) was purchased from Sigma-Aldrich (St. Louis, MO). Adjuplex (ADJ), CpG-ODN 1826 (CpG) oligonucleotide and Glucopyranosyl Lipid Adjuvant (GLA) were purchased from Empirion LLC (Columbus, OH), InvivoGen (San Diego, CA) and Avanti Polar Lipids, Inc. (Alabaster, AL), respectively. For footpad immunization, mice were briefly anesthetized with isoflurane, after which 15 ug of Alexa Fluor 647-conjugated chicken OVA (Thermo Fisher) formulated in saline, ADJ (5%), CpG (5ug), GLA (5ug), ADJ (5%)+CpG (5ug) or ADJ (5%) + GLA (5ug) was injected to hind footpad. For SQ vaccination, C57BL/6J mice were vaccinated at the tail base with 50ul of the vaccine: ovalbumin (10ug) formulated in saline, ADJ (5%), CpG (5ug), GLA (5ug), ADJ (5%) + CpG (5ug) or ADJ (5%) + GLA (5ug). Mice were boosted after 21 days of the initial vaccination.

### Tissue processing and Flow cytometry

The vaccine-draining popliteal LN was incubated with 3ml of filtered 1mg/ml collagenase D (Roche) for 30 minutes at 37 C’. The digested lymph nodes were further mechanically processed, filtered, to generate single cell suspensions. Spleens were mechanically processed into single-cell suspensions using standard procedures. Single-cell suspensions were first stained for viability with LiveDead eFlour 780 stain (eBioscience) for 30 minutes on ice. Next, samples were stained with antibodies diluted in Brilliant Stain Buffer (BD Biosciences) for 30 minutes (for innate immune cells, as previously described (41)) or 60 minutes (for T cell immunophenotyping) with K^b^/SIINFEKL tetramers (provided by Emory MHC tetramer facility, Atlata) and antibodies listed in Supplementary table 1. Following staining, cells were washed with FACS buffer (0.1% BSA + PBS) twice and fixed with 2% paraformaldehyde for 10 minutes on ice. Samples were washed again twice with FACS buffer and acquired on LSRFortessa (BD Biosciences) and analyzed with FlowJo V.10 software (TreeStar, Ashland, OR).

### Intracellular staining for transcription factors

To stain for transcription factors or granzyme B, cells were first stained for viability with LiveDead eFlour 780 stain (eBioscience) for 30 minutes and then stained with antibodies and MHC I tetramers diluted in Brilliant Stain Buffer (BD Biosciences) for 60 minutes. The samples were then fixed, permeabilized and subsequently stained for transcription factors using the transcription factors staining kit (eBioscience) with the antibodies listed in Supplementary table 1 in Perm Wash buffer. All samples were acquired on LSRFortessa (BD Biosciences) and analyzed with FlowJo V.10 software (TreeStar, Ashland, OR).

### Intracellular cytokine staining

For intracellular cytokine staining, one million cells (1×10^6^) cells were plated on flat-bottom tissue-culture-treated 96-well plates (Corning) Cells were stimulated for 5 hours at 37C in the presence of brefeldin A (1 μl/ml, GolgiPlug, BD Biosciences), human recombinant IL-2 (10 U/well) and with or without SIINFEKL peptide (Genscript) at 0.2ug/ml. After *ex vivo* peptide stimulation, cells were stained for viability dye (LiveDead eFluor 780) for 30 minutes, stained with surface antibodies, and fixed/permeabilizated with Cytofix/Cytoperm kit (BD Biosciences, Franklin Lakes, NJ) according to manufacturer’s protocol. Samples were stained with anti-cytokine antibodies listed in Supplementary table 1 in perm wash buffer for 30 minutes, washed with perm wash buffer, and re-suspended in FACS buffer before flow cytometry.

### Vaccination and enumeration of Listeria Challenge

At > 40 days after booster vaccination, mice were challenged intravenously with 1.7 x 10^5^ CFUs of LM-OVA (*Listeria monocytogenes* expressing chicken ovalbumin (LM-OVA), provided by Dr. Hao Shen (University of Pennsylvania School of Medicine). To quantify Listeria burdens, tissues were homogenized in GentleMACS C-Tubes (Miltenyi) via GentleMACS dissociator. Organs were processed in sterile 0.1% Nonidet-P40 (VWR) + PBS in gentleMACS C Tubes. Serial dilutions of tissue samples (undiluted to 10^6^) were plated on brain heart infusion agar plates for 24 hours at 37 C’. Listeria burden in tissues were normalized by the weight of the tissues.

### Statistical analyses

Statistical analyses were performed using GraphPad software 8.1.1 (La Jolla, CA). All comparisons were made using either Mann-Whitney U test or an one-way ANOVA test with Tukey corrected multiple comparisons where p<0.05 = *, p<0.005 = **, p<0.0005 = *** were considered significantly different among groups. Bacterial titers were log transformed prior to analysis. One statistical outlier was excluded from analysis of CpG-vaccinated mice in Listeria challenge experiment.

**Figure 5.**
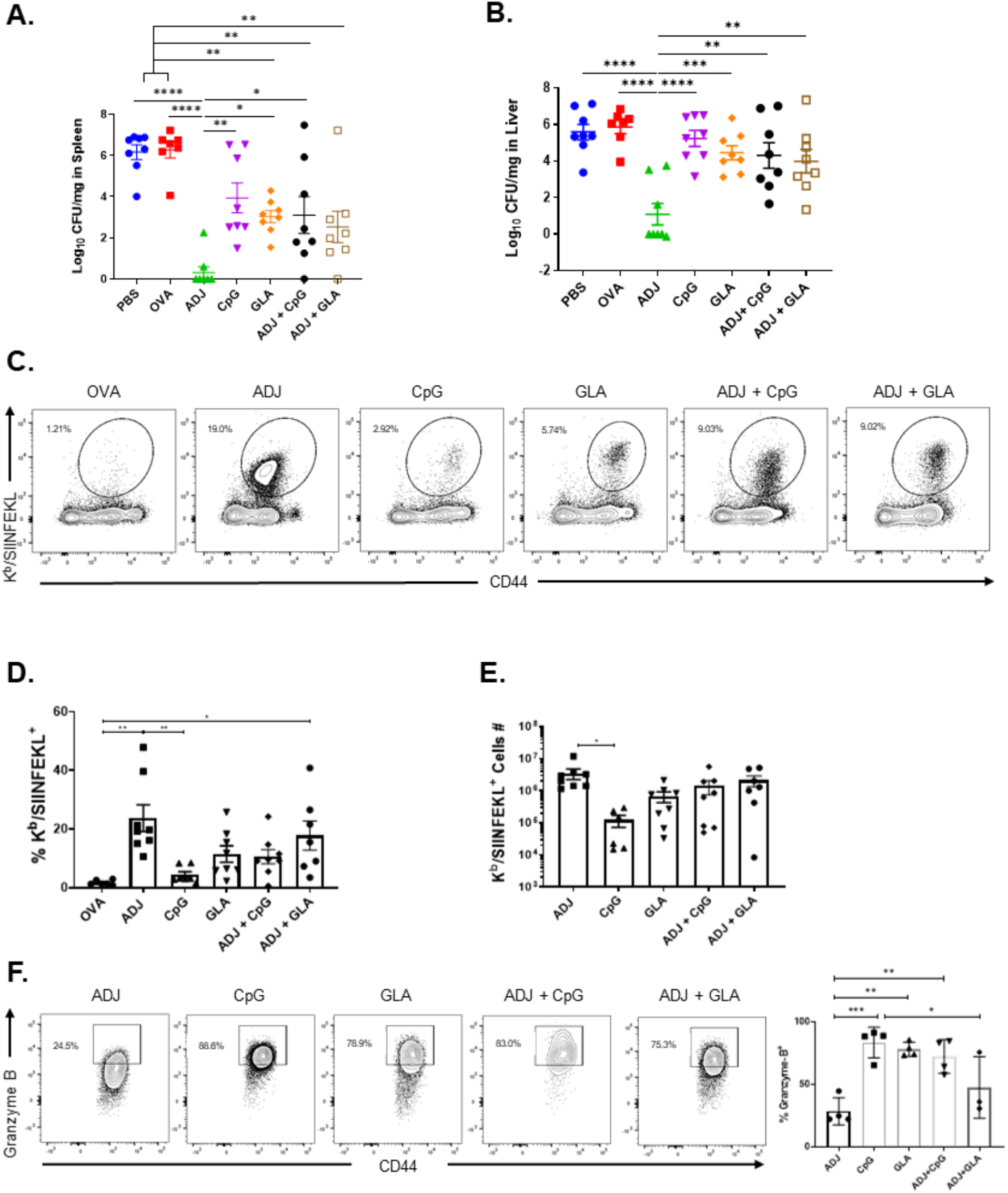
Vaccine-induced protective immunity to Listeria. Mice were vaccinated twice (3 weeks apart) SQ with OVA (10ug/mouse) formulated in ADJ, CpG, GLA, ADJ+CpG, ADJ+GLA. Forty days after the booster vaccination, mice were challenged with virulent recombinant OVA-expressing Listeria (LM-OVA) and sacrificed at day 5. (A) Listeria burden was quantified in the spleens and livers on the 5^th^ day after challenge. (B) Single-cell suspensions of spleen were stained with D^b^/SIINFEKL tetramers, anti-CD8, anti-CD44, and anti-granzyme B antibodies. (C) or (F) FACS plots show percentages of gated tetramer-binding CD8 T cells in respective gates. (D-E) Percentages and total numbers of OVA SIINFEKL-specific CD8 T cells in spleens. Data in each graph show Mean ± SEM. Data are pooled from two independent experiments (A-E) or represent one of two independent experiments (F). Each independent experiment had n=3-5 mice per group. *, **, ***, and **** indicate significance at *P*<0.1, 0.01, 0.001, and 0.0001 respectively. (One-way ANOVA: A-F)

## Acknowledgements

We would like to thank all the members of Suresh Laboratory for constructive feedback and technical assistance and genuine appreciation for the efforts of the veterinary and animal care staff at UW-Madison. Thanks to Zachary Morrow (Dr. JD Sauer Lab) for Listeria challenge experiment. We are thankful to the Emory NIH Tetramer Core Facility for providing MHC-I tetramers.

## Author contributions

WL, AL and MS designed, performed, analyzed experiments, and provided conceptual input for the manuscript. BB provided technical expertise and intellectual insights for the manuscript. WL and MS wrote the manuscript, which was proofread by all authors.

---

**Supplementary Figure 1.**
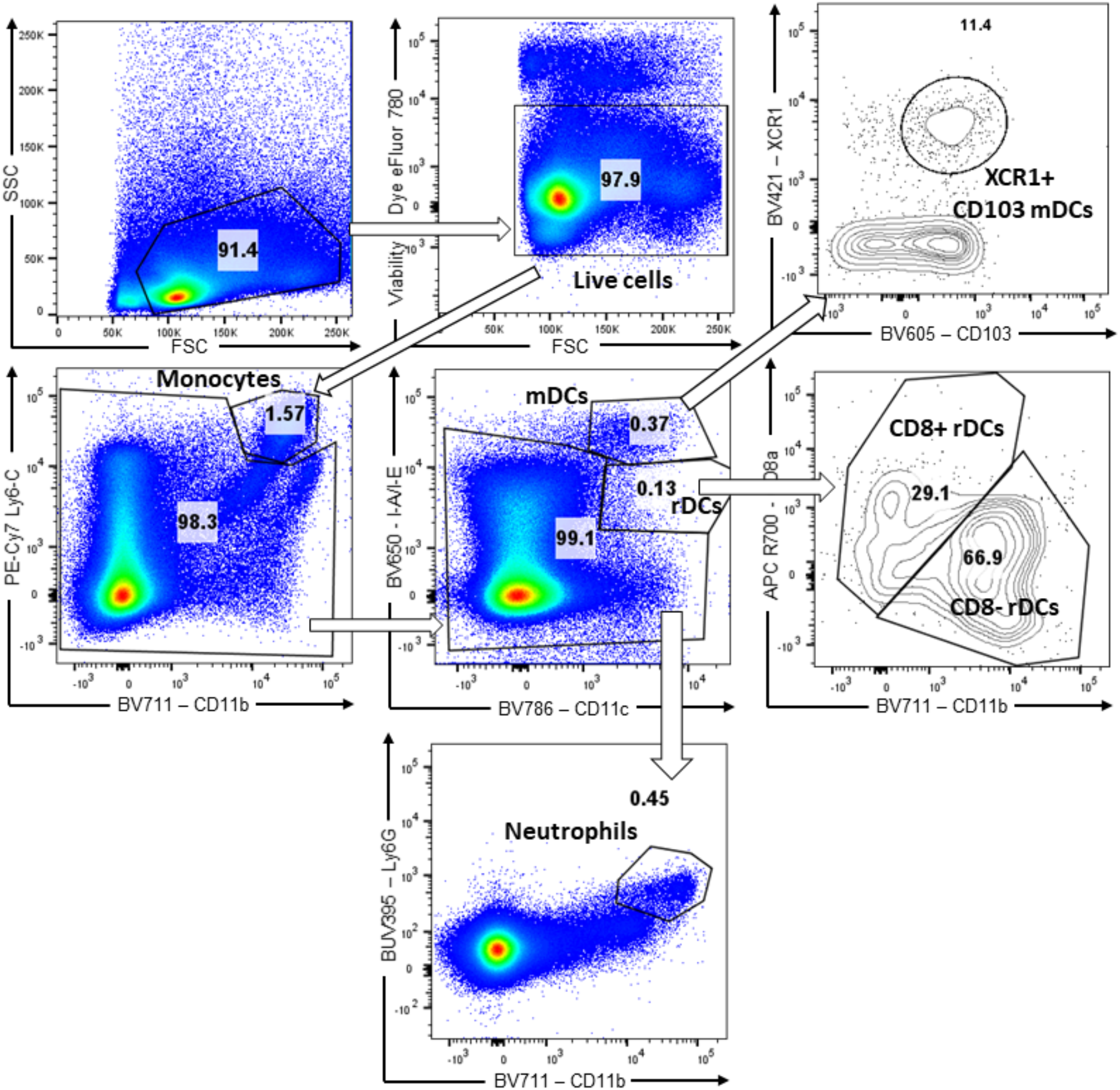
Analysis of innate immune cells in draining nodes of vaccinated mice. Gating strategy for innate immune cell subsets in draining popliteal lymph nodes (DLNs). Single cell suspensions of DLN were stained with antibodies to Ly6G, XCR1, CD80, CD86, CD103, MHC-II, CD11b, CD11c, Ly6C, and CD8α.

**Supplementary Figure 2.**
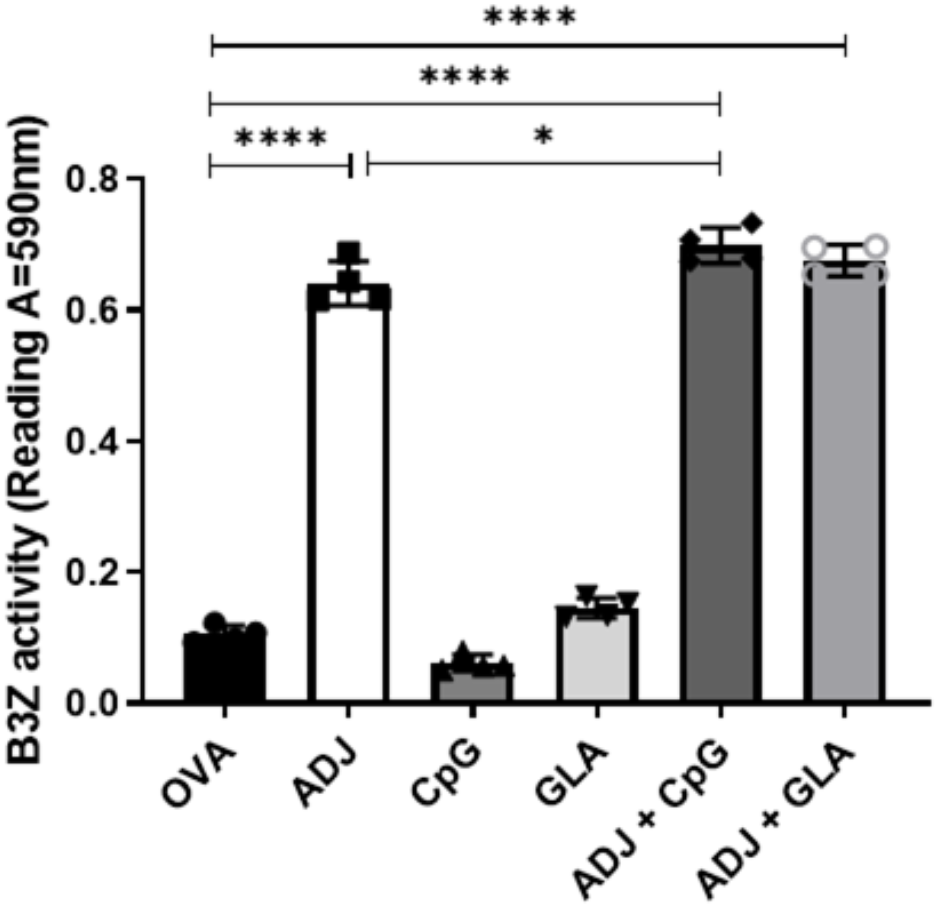
Effect of combination adjuvants on cross-presentation by bone marrow-derived DCs. BMDCs were exposed to media or OVA ± ADJ, CpG or GLA for 5 h, and co-cultured with B3Z cells for 24 h. β-galactosidase in co-cultured B3Z cells was quantified by CPRG colorimetry. Data in graph show Mean ± SEM. Data represent one of two independent experiments. Each independent experiment had triplicates per group. *, **, ***, and **** indicate significance at *P*<0.1, 0.01, 0.001, and 0.0001 respectively.

**Supplementary Table 1:** List of antibodies and tetramers used in this study.

| **Antibody (Dilution Factor)** | **Company** | **Catalogue number** |
| --- | --- | --- |
| Rat anti-mouse Ly-6G-conjugated (1:200) | BD Biosciences | 563978 |
| Rat anti-mouse I-A/I-E-BV650-conjugated (1:400) | BD Biosciences | 563415 |
| Rat anti-mouse CD11b-BV711-conjugated (1:400) | BD Biosciences | 563168 |
| Hamster anti-mouse CD11c-BV786-conjugated (1:100) | BD Biosciences | 563735 |
| Rat anti-mouse CD86-BV510-conjugated (1:200) | BD Biosciences | 563077 |
| Rat anti-mouse Ly-6C-PE-Cy7-conjugated (1:400) | BD Biosciences | 560593I |
| Rat anti-mouse CD8 α-APC-R700-conjugated (1:200) | BD Biosciences | 564983 |
| Rat anti-mouse CD8α-BUV395-conjugated (1:200) | BD Biosciences | 563786 |
| Rat anti-mouse CD4-BUV496-conjugated (1:200), | BD Biosciences | 564667 |
| Rat anti-mouse CD44-BV510-conjugated (1:200) | BD Biosciences | 563114 |
| Rat anti-mouse CD62L-PE-CF594-conjugated (1:200) | BD Biosciences | 562404 |
| Rat anti-mouse CD62L-PE-CF594-conjugated (1:200) | BD Biosciences | 562404 |
| Rat anti-mouse B220-PerCP-conjugated (1:200) | BD Biosciences | 552879 |
| Hamster anti-mouse-CD69-PE-Cy7-conjugated (1:200) | BD Biosciences | 553237 |
| BUV496-streptavidin (1:400) | BD Biosciences | 564666 |
| mouse anti-mouse T-bet-BV421 conjugated (1:200) | BD Biosciences | 563318 |
| rat anti-mouse IFN-γ-APC-conjugated (1:400) | BD Biosciences | 554413 |
| rat anti-mouse TNF-α-BV421-conjugated (1:500) | BD Biosciences | 554413 |
| rat anti-mouse XCR1-BV421-conjugated (1:100) | Biolegend | 148216 |
| rat anti-mouse CD103-BV605-conjugated (1:200) | Biolegend | 121433 |
| mouse anti-mouse CD64 (FcγRI) Antibody (1:200) | Biolegend | 139308 |
| rat anti-mouse CD127- PerCP/Cyanine5.5 (1:150) | Biolegend | 135022 |
| hamster anti-mouse KLRG1-BV711-conjugated (1:200) | Biolegend | 564014 |
| mouse anti-mouse CX3CR1-BV785-conjugated (1:200) | Biolegend | 149029 |
| Hamster anti-mouse-CD80-biotin-conjugated (1:200) | Biolegend | 104704 |
| Rat anti-mouse Granzyme B-BV421-conjugated (5ul/well). | Biolegend | 396414 |
| Rat anti-mouse CD19-PE-Cy7-conjugated (1:200) | eBioscience | 25-0193-82 |
| Rat anti-mouse Granzyme B-PE-conjugated (5ul/well) | eBioscience | MHGB04 |
| rat anti-mouse EOMES-PE-eFluor 610 conjugated (1:200) | eBioscience | 61-4875-82 |
| mouse anti-human BATF-PerCP-eFluor 710 conjugated (5ul/well) | eBioscience | 46-9860-42D |
| APC-conjugated H2-K<sup>b</sup> tetramers bearing the SIINFEKL peptide (1:150) | NIH Tetramer Core Facility | N/A |
| Purified Anti-Mouse CD16 / CD32 (Fc Shield) (2.4G2) | Tonbo Biosciences | 70-0161-U500 |

